# Dissecting disgust and fear in chimpanzees: From facilitation to disruption of cognitive processes

**DOI:** 10.1101/2025.11.20.689418

**Authors:** Cécile Sarabian, Andrew J.J. MacIntosh, André Gonçalves, Nobuyuki Kawai, Ikuma Adachi

## Abstract

Disgust and fear are defensive emotions that evolved to address different types of threat. Disgust reduces pathogen exposure through avoidance, whereas fear promotes vigilance and escape from predators. While cognitive mechanisms of fear have been extensively studied, those underlying disgust remain less understood, particularly in nonhuman animals. We examined how cues of disease and predation risk influence cognitive processes in captive chimpanzees (*Pan troglodytes*). In three touchscreen experiments, subjects performed a number ordering task while exposed to disgust-related images (carcasses, rotten food, disease-associated invertebrates), pathogen-related odors (cadaverine, butyric acid) or fear-related images (snakes). Task performance was assessed via accuracy (proportion of correct trials), reactivity (latency to touch the first numeral), and efficiency (latency to complete trials). We found that disgust cues – particularly those related to body decomposition and food spoilage – modulated task performance in distinct temporal patterns. Carcass images reduced accuracy relative to invertebrate images and slowed performance compared to snake images, while butyric acid – and to a lower extent cadaverine – initially facilitated but later disrupted efficiency upon re-exposure. In contrast, snake images enhanced performance by reducing latencies. To test whether these effects reflected differences in visual attention, we conducted a fourth eye-tracking experiment with new carcass and snake images. Both types of stimuli captured and held attention, partly echoing human data showing sustained attention to snakes but typically not to disgust images. Overall, our results reveal distinct cognitive signatures of disgust and fear in chimpanzees, offering comparative insight into the evolution of defensive emotions.

**Significance statement:** Disgust and fear are fundamental emotions that evolved to protect organisms from distinct kinds of threats – disease and predation – but their effects on cognition have rarely been compared, especially in nonhuman animals. In this study, we examined how chimpanzees respond to pathogen-and predator-related cues using touchscreen and eye-tracking experiments. We found that pathogen cues influenced cognitive performance in complex, time-dependent ways, sometimes enhancing and later disrupting it, while predator cues consistently improved performance. Both types of stimuli captured and held attention, although sustained attention to pathogen cues was unexpected. These findings reveal distinct emotional and cognitive signatures of disgust and fear, shedding light on the evolutionary roots of how emotions shape behavior and attention.

## INTRODUCTION

Disgust and fear are two emotions that evolved to serve different adaptive functions. While disgust primarily protects from disease by motivating the avoidance of pathogens, parasites and toxins (*1–3*), fear facilitates escape from immediate threats, such as predators, via fight-or-flight responses (*4–7*). The differing consequences of these threats – from acute or chronic disease and reduced fitness in the case of infection to immediate death in the case of predation – may have led to the evolution of distinct responses. In humans, disgust is expressed by facial expressions that block the entry of potential toxins or pathogens (narrowed eyes, nose wrinkling, raised upper lip) (*8–10*), oculomotor avoidance after initial processing (*11–13*), and physiological reactions such as pupil constriction (*14, 15*) – or dilation (*16*) – an inconsistency, which could be due to differences in stimulus type. Disgust also disrupts gastric rhythm (*17, 18*), reduces heart rate (*19, 20*) and activates the anterior insula (*21, 22*). By contrast, fear manifests through expressions that enhance sensory perception (raised eyebrows, nose elongation, stretched lips) (*9*), early and sustained attention to the threat (*23, 24*), along with pupil dilation (*16,25*), elevated heart rate (*26*), adrenaline release (*27*), and amygdala activation (*28*). Disparities in perception and outcomes may have contributed to the research gap between these two emotions, with fear historically receiving more attention, though interest in disgust has markedly increased since the 2000s (*29, 30*). Disgust has been understudied not only in humans but across species (*31*). Moreover, these two emotions are frequently examined in isolation, even though threats from pathogens and predators often co-occur within the same environments (*30, 32*). This siloed approach limits the ecological validity of current findings. Understanding how disgust and fear interact – or diverge – is crucial for uncovering the evolutionary mechanisms underlying threat detection and behavioral responses, particularly in our closest phylogenetic relatives, where emotional functioning remains largely unexplored.

Recent research suggests that non-human primates display core disgust responses similar to those observed in humans, including characteristic facial expressions (*33–35*), physiological reactions like nausea (*34*), anterior insula activation (*33, 34*), and avoidance behaviors (*36–43*). Despite increasing evidence of emotional expressions and physiological responses associated with disgust in non-human primates, its cognitive dimension – how it modulates attention and decision-making – has received little empirical investigation. In contrast, fear has been extensively studied behaviorally, cognitively and physiologically (*44, 45*). Notably, both emotions have been primarily examined through visual stimuli in primates, e.g. (*46, 47*), while threat perception may involve sensory diversity (*48–50*).

In this study, we explored the effect of pathogen-versus predator-related cues on cognitive processes in captive trichromatic chimpanzees (*Pan troglodytes*), building on previous findings of chimpanzees’ avoidance behaviors toward pathogen-associated cues (*36, 51*). We conducted four experiments to address the following questions: 1) Do disgust-related images influence cognitive performance, here defined as the accuracy and latency with which individuals process and respond to task stimuli? 2) Do pathogen-associated odors influence cognitive performance? 3) Does cognitive performance differ under disgust-versus fear-related stimuli? 4) Do attentional patterns differ between disgust- and fear-related stimuli? To test these, chimpanzees completed a *Number ordering task* (*52*), a standard touchscreen test of working memory and attention, while being presented with visual or olfactory stimuli. Attentional responses were measured using eye-tracking. In line with the hypothesis that disgust promotes the avoidance of physical and perceptual contact with pathogen sources (*53–55*), we predicted that pathogen-related cues would impair cognitive performance (e.g. reduce accuracy, reactivity and/or efficiency), while fear-related cues would enhance it due to heightened vigilance (e.g. (*56*)). According to the eye-mind hypothesis – that gaze reflects attention and thought (*57*) – we also expected that disgust images would divert gaze and reduce attentional focus, while fear images would sustain visual attention. Understanding how these emotions modulate cognition offers insight into their adaptive roles in guiding behavior under threat.

## RESULTS

### Disgust-related images selectively affect performance

We began by testing whether and which disgust-related images (DIRTI (*58*), see SI Appendix, Fig. S1-S3) influence chimpanzee performance on a number ordering task – where numerals from 1 to 9 were presented simultaneously at random locations on a touchscreen and had to be selected in ascending order (Fig. 1A). Every five trials, a full-screen image was displayed: either a disgust-related one – such as a heterospecific animal carcass (mammal, bird, amphibian or marsupial, at various stages of decay), rotten food (spoiled fruits or vegetables), invertebrates associated with disease risk (e.g. tick, mosquito) – or a corresponding control image (a sleeping animal, fresh food, butterflies or other non-pathogenic invertebrates). Given that the three control conditions did not significantly differ in the proportion of correct trials (χ^2^(2) = 2.20, *p* = 0.333), they were combined into a single control category for subsequent analyses. Performance was measured via trial accuracy (correct vs. incorrect), reactivity (latency to touch the first numeral), and time to complete correct trials.

**Fig. 1:**
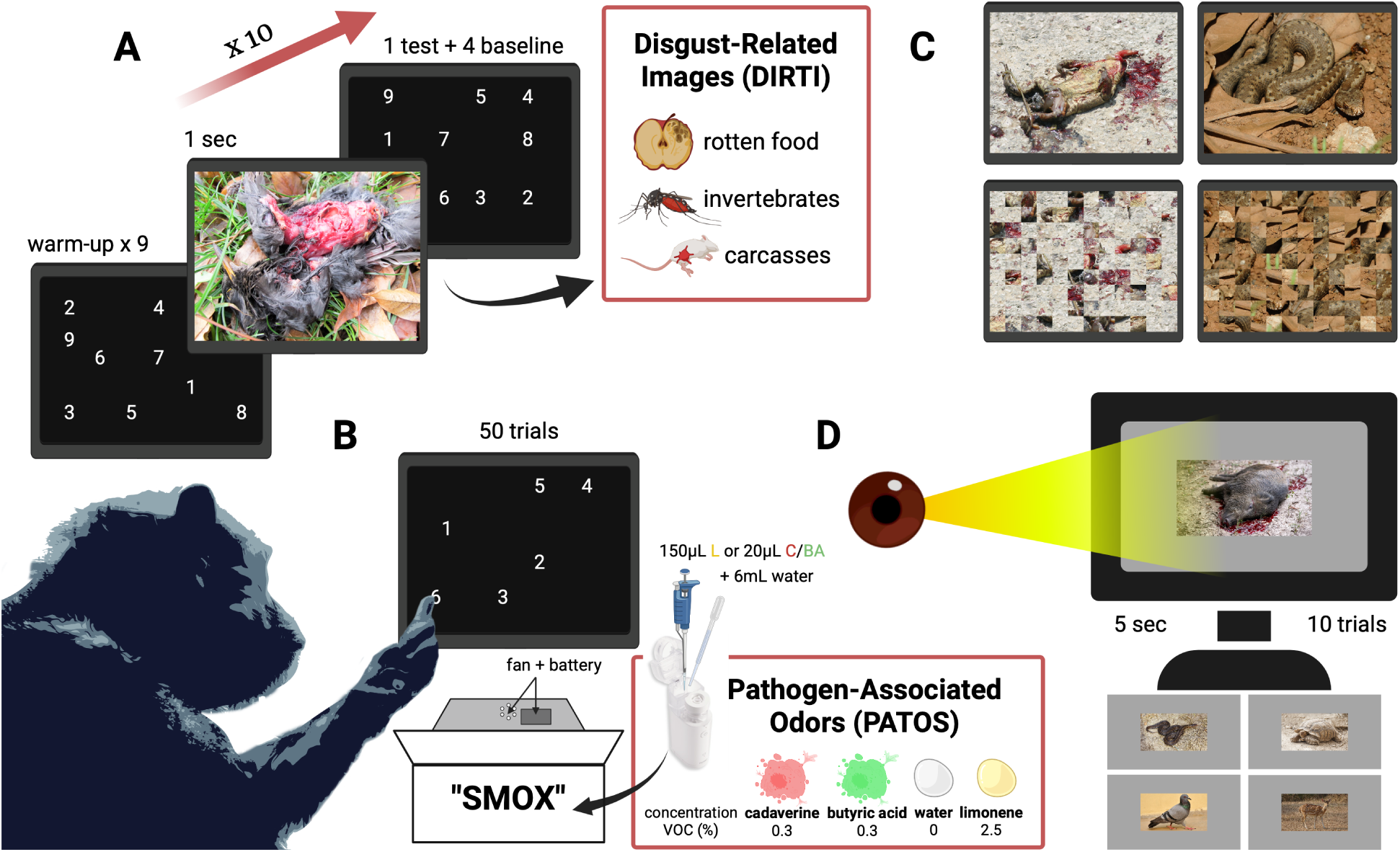
Experimental designs of number ordering (A-C) and eye-tracking (D) tasks. (**A**) Experiment 1: Disgust-related images of rotten food (i.e. spoiled fruits and vegetables), disease-associated (e.g. mosquitoes)/pathogen-mimicking (e.g. earthworms) invertebrates, heterospecific animal carcasses (from birds, mammals, amphibians and marsupials) or their controls were presented for one second at regular intervals on touch screens during a number ordering task. Chimpanzees had to touch numerals from one to nine in an ascending order to complete a trial and performed a total of three sessions of 50 trials per condition. (**B**) Experiment 2: Pathogen-associated odors of cadaverine, butyric acid or their controls were diffused at low concentrations via a self-designed odor box (*SMOX*) while chimpanzees performed the number ordering task. Numerals were reduced from nine to six to ease completion and reduce potential discomfort. (**C**) Experiment 3: New heterospecific animal carcass, snake or control scrambled images of those were presented at regular intervals on touch screens during a number ordering task using the same paradigm as in A. (**D**) Experiment 4: New sets of animal carcass and snake images or their controls (i.e. deer, pigeon, tortoise) were presented for five seconds while chimpanzees sucked juice through the plexiglass to maintain their head straight. Gaze patterns on each image were recorded via an eye-tracker. Figure created with BioRender. Artwork from original chimpanzee (Ai) image: Kenneth Keuk.

The generalized linear mixed model (GLMM) for accuracy (M1.1) including the interaction between condition and trial number did not significantly improve model fit over the additive model without the interaction (LRT; Δ*LogLik* = 3.6, Δ*d*. *f*. = 3, *p* = 0.062), but the additive model outperformed the null model (Δ*LogLik* = 4.5, Δ*d*. *f*. = 3, *p* = 0.031, see SI Appendix Table S2). We therefore retained the additive model. The full model for reactivity (M1.2), however, did not outperform its null model (Δ*LogLik* = 5.1, Δ*d*. *f*. = 6, *p* = 0.114) and was therefore not retained. In contrast, for latency to complete correct trials (M1.3), the model including the interaction between condition and number of image exposures significantly outperformed both its null (Δ*LogLik* = 9.4, Δ*d*. *f*. = 6, *p* = 0.004) and additive (Δ*LogLik* = 7, Δ*d*. *f*. = 3, *p* = 0.003) models and was retained for interpreting effects of image condition.

The proportion of correct trials was high across conditions (*invertebrate*: 0.85, *control*: 0.84, *rotten food*: 0.83, *carcass*: 0.80; Fig. 2A). However, the GLMM revealed significant differences in accuracy across image conditions (χ^2^(3) = 8.9, *p* = 0.031; Table S2). Accuracy was significantly lower in the *carcass* condition compared to the *control* (β = –0.24 ± 0.10 SE, *z* = –2.44, *p* = 0.015), while differences for the *invertebrate* (β = 0.11 ± 0.11 SE, *z* = 0.96, *p* = 0.335) and *rotten food* (β = −0.10 ± 0.10 SE, *z* = −0.93, *p* = 0.354) conditions were not significant (Table S3). Trial number itself did not influence accuracy (β = 0.04 ± 0.04 SE, *z* = 0.99, *p* = 0.323). The estimated marginal means confirmed this pattern: *invertebrate* = 1.80 [1.44, 2.16], *control* = 1.69 [95% CI: 1.37, 2.02], *rotten food* = 1.60 [1.24, 1.95], and *carcass* = 1.45 [1.10, 1.80]. Post hoc pairwise comparisons indicated that the effect was mainly driven by the difference between *carcass* and *invertebrate* conditions (β = −0.35 ± 0.13 SE, *z* = −2.74, *p* = 0.031; Table S4).

**Fig. 2:**
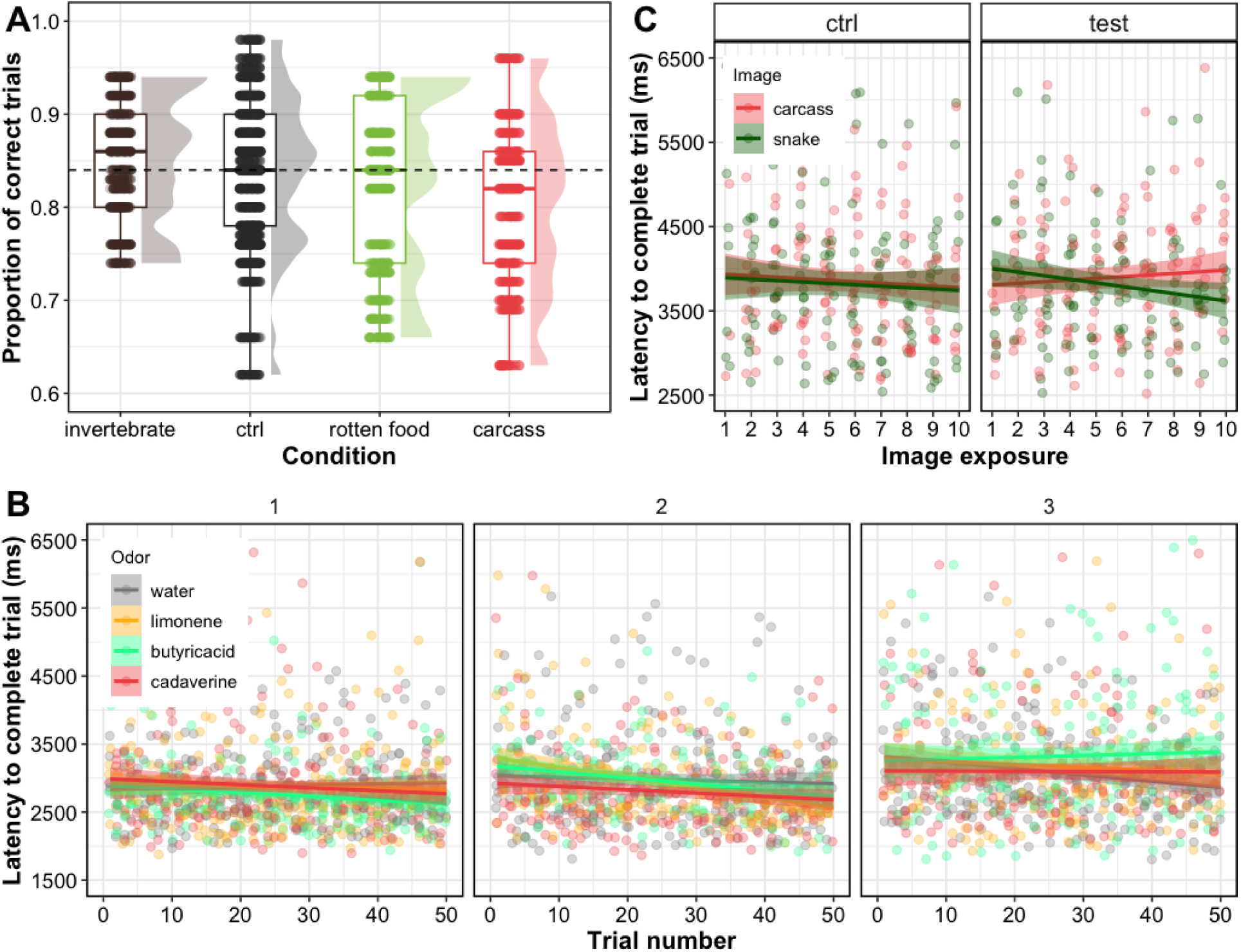
Effect of disgust- and fear-related stimuli on chimpanzee cognitive performance in the number ordering task. (**A**) Experiment 1: Raincloud plots show the proportion of correct trials during sessions with images of pathogen-associated/mimicking invertebrates (brown), controls (black), rotten food (green), and animal carcasses (red). Chimpanzees made significantly more errors when exposed to carcass images compared to invertebrate images (*p* < 0.05), but not compared to rotten food or control images. (**B**) Experiment 2: Scatter plots display the latency to complete correct trials (in ms) under exposure to water (grey), limonene (yellow), butyric acid (green), and cadaverine (red) odors in sessions 1-3. Chimpanzees were significantly slower under repeated exposure to butyric acid than under water (*p* < 0.05). (**C**) Experiment 3: Scatter plots of latency (ms) in sessions with carcass (red) and snake (green) images. Chimpanzees were significantly faster under repeated exposure to snake images compared to carcass images (*p* < 0.01).

Task efficiency, measured by the latency to complete trials, significantly improved with exposure to *rotten food* images compared to the *control* (β = −0.04 ± 0.01 SE, *z* = −3.13, *p* = 0.002; Table S3), whereas other conditions did not differ from the control (*invertebrate*: β = 0.00 ± 0.01 SE, *z* = 0.23, *p* = 0.816; *carcass*: β = 0.02 ± 0.01 SE, *z* = 1.16, *p* = 0.247). Post hoc pairwise comparisons showed that chimpanzees were faster after the first few *rotten food* images (versus *control*: β = 0.10 ± 0.03 SE, *z* = 3.34, *p* = 0.005; versus *invertebrate*: β = 0.12 ± 0.03 SE, *z* = 3.53, *p* = 0.002; versus *carcass*: β = 0.13 ± 0.03 SE, *z* = 3.69, *p* = 0.001), but that effect did not persist after multiple exposures (see Table S4).

### Pathogen-associated odors modulate efficiency over time

Building on the effects of disgust-related images, we next tested whether pathogen-associated odors similarly affect cognitive performance. In this second experiment, we used the same paradigm as before, but replaced the visual stimuli with olfactory ones. We diffused low concentrations (see Materials and Methods) of different odors – cadaverine (associated with putrefaction), butyric acid (vomit-like odor), limonene (citrus scent) or water – through a box and nebulizer built just beneath the touchscreen, while chimpanzees performed the task (Fig. 1B; Movie S1). To shorten the sessions and minimize potential discomfort, the number of numerals presented on the screen was reduced from nine to six.

Only the GLMM for latency to complete correct trials (M2.3) showed a significantly better fit when including the interactions between odor, trial and session numbers compared to both the null (LRT; Δ*LogLik* = 16, Δ*d*. *f*. = 13, *p* = 0.001) and the additive (without interactions; Δ*LogLik* = 15, Δ*d*. *f*. = 10, *p* < 0.001) models (Table S2). We therefore retained this interaction model for interpretation.

Overall, latency to complete the task increased across trials and sessions under both *cadaverine* (GLMM; β = 0.06 ± 0.02 SE, *z* = 3.60, *p* < 0.001) and *butyric acid* (β = 0.05 ± 0.02 SE, *z* = 3.12, *p* = 0.001) relative to the control *water* condition, whereas the effect of *limonene* was not significant (β = 0.02 ± 0.02 SE, *z* = 1.40, *p* = 0.160; Fig. 2B; Table S5). Post hoc pairwise comparisons revealed more complex temporal dynamics. During the last third of the first session (after ≈ 4 min of diffusion), chimpanzees were faster under *cadaverine* (β = −0.12 ± 0.05 SE, *z* = –2.57, *p* < 0.05) and *butyric acid* (β = −0.14 ± 0.05 SE, *z* = –2.75, *p* = 0.031) compared to *water* (Table S6). This effect reversed during the first two-thirds of the second session, though differences were not significant. In the middle to late portion of the third session, *butyric acid* slowed latency relative to water (*mid*: β = 0.11 ± 0.04 SE, *z* = 3.04, *p* = 0.013; *high*: β = 0.16 ± 0.05 SE, *z* = 3.12, *p* < 0.01), whereas *cadaverine* showed a similar but non-significant trend (*mid*: β = 0.04 ± 0.04 SE, *z* = 1.07, *p* = 0.708; *high*: β = 0.12 ± 0.05 SE, *z* = 2.29, *p* = 0.102). Although *limonene* did not affect overall latency, early trials in the second session were slower compared to *water* (β = 0.14 ± 0.05 SE, *z* = 2.70, *p* = 0.035), consistent with the direction observed for *cadaverine* and *butyric* acid.

### Carcass and snake images have opposite effects on task efficiency over time

In a third experiment, we used new sets of animal carcass images – identified in Experiment 1 as eliciting the lowest proportion of correct trials – and compared their effect to snake images, using the same paradigm. We used scrambled versions of the carcass and snake images as controls (Fig. 1C).

Only the GLMM for latency to complete correct trials (M3.3) with interactions between condition, image type and number of exposures outperformed its respective null (LRT; Δ*LogLik* = 7.1, Δ*d*. *f*. = 6, *p* = 0.027) and additive (without the interaction; Δ*LogLik* = 6.9, Δ*d*. *f*. = 4, *p* = 0.008) models (Table S2).

Chimpanzees completed the task significantly faster when exposed to *snake* images compared to *carcass* (GLMM; β = −0.06 ± 0.02 SE, *z* = –2.59, *p* < 0.001) images (Fig. 2C; Table S7; Movies S2, S3). Pairwise post hoc comparisons confirmed this effect, showing that repeated exposure to *snake* images improved performance (β = −0.10±0.03 SE, *z* = –3.17, *p* = 0.006), whereas exposure to *carcass* images slowed it down. Other comparisons at minimum, mid and high image exposure were not significant (all *p* > 0.090; see Table S8).

### Carcass and snake images attract and retain attention

In the fourth experiment, we used an eye-tracking task to present chimpanzees with new sets of animal carcass and snake images, as well as control images of tortoises, pigeons and deer. This allowed us to measure their visual attention and test whether attentional patterns could explain the performance differences observed in Experiment 3. We used three proxies for visual attention: the location of the first fixation (on the animal or not), the proportion of fixations on the animal versus elsewhere (i.e. the environment in the picture or the grey area surrounding it; Fig. 1D; see Fig. S4) and the proportion of time spent looking at the animal versus looking elsewhere (hereafter *attention score*).

All three GLMMs (M4.1-3) with their main effect were significantly better than their null, intercept-only, models (LRT; first fixation location: Δ*LogLik* = 14.9, Δ*d*. *f*. = 4, *p* < 0.001; proportion of fixations: Δ*LogLik* = 32.7, Δ*d*. *f*. = 4, *p* < 0.001; proportion of time looking: Δ*LogLik* = 42.7, Δ*d*. *f*. = 4, *p* < 0.001; Table S2).

### First fixation location

Chimpanzees’ first fixation was significantly more likely to land on the animal in *carcass* images compared to control images (GLMM; all *p* < 0.01; Fig. 3A; Table S9). However, there was no significant difference in the likelihood of first fixating on the animal in *carcass* versus *snake* images (β < −0.01 ± 0.24 SE, *z* = –0.01, *p* = 0.995). Pairwise post hoc comparisons indicated that this effect was mainly driven by contrasts with *pigeon* images (*carcass*–*pigeon*: β = 1.39 ± 0.30 SE, *z* = 4.66, *p* < 0.001; *snake*–*pigeon*: β = 1.39 ± 0.30 SE, *z* = 4.58, *p* < 0.001), whereas all other comparisons were non-significant (*p* > 0.06; Table S10).

**Fig. 3:**
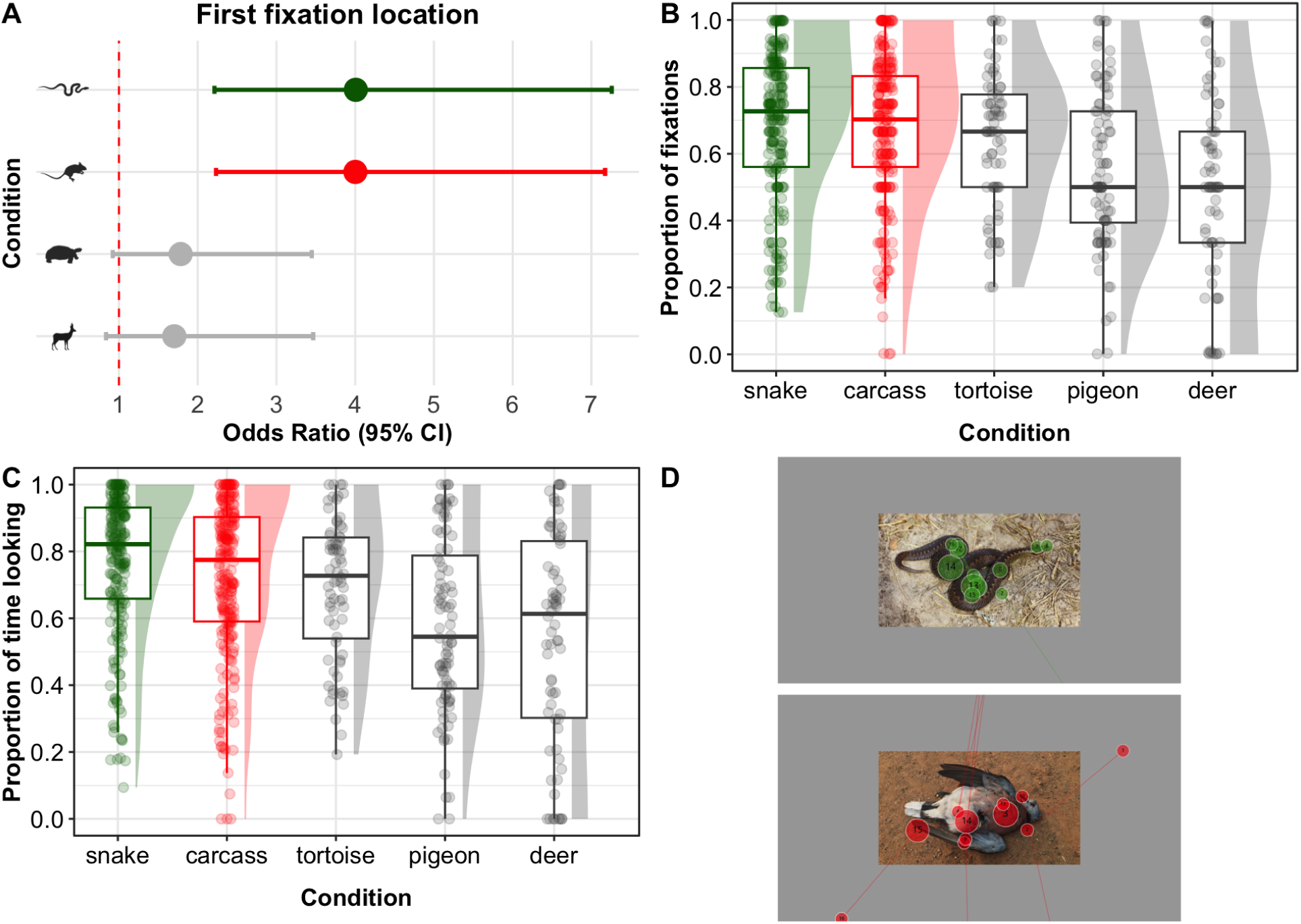
Effect of disgust- and fear-related images on patterns of visual attention in Experiment **4.** (**A**) Forest plots show the odds ratios of the first fixation location on snakes (green), carcasses (red), tortoises (grey) and deer (grey), taking the pigeon images as referential. Chimpanzees’ first fixation had significantly more chances to land on the snake or carcass than on the pigeon (both *p* < 0.001). (**B**) Raincloud plots for the proportion of fixations on snake (green), carcass (red), tortoise, pigeon and deer (grey) images. Chimpanzees had more fixations on snakes and carcasses compared to deer and pigeons (all *p* < 0.001). (**C**) Raincloud plots for the proportion of time looking at the snake, carcass, tortoise, pigeon and deer in the image vs. looking away (i.e. attention score). Chimpanzees looked at carcasses and snakes for longer than at control animals (all *p* < 0.01). However, there was no significant differences between snakes and carcasses (*p* = 0.210). (**D**) Gaze plots of chimpanzees looking at a snake (Ayumu, session 1, slide 1; top) and a bird carcass (Ai, session 1, slide 6; bottom). Fixations are shown as circles, with size proportional to fixation duration. Lines between circles indicate saccades – rapid eye movements during which vision is suppressed.

### Proportion of fixations

Chimpanzees directed a significantly greater proportion of fixations toward the animal in *carcass* images than in control images (GLMM; all *p* < 0.05, Fig. 3B; Table S9), whereas no difference was found relative to *snake* images (β = −0.03 ± 0.06 SE, *z* = –0.53, *p* = 0.599). Pairwise post hoc comparisons revealed that animals in *carcass* images received more fixations than those in *deer* (β = 0.66 ± 0.10 SE, *z* = 6.90, *p* < 0.001) and *pigeon* (β = 0.45 ± 0.09 SE, *z* = 5.05, *p* < 0.001) images, but not more than in *snake* (β = −0.03 ± 0.06 SE, *z* = –0.53, *p* = 1.000) or *tortoise* (β = 0.19 ± 0.10 SE, *z* = 2.01, *p* = 0.260; Table S10) images. Similarly, animals in *snake* images attracted more fixations than those in *deer* (β = 0.69 ± 0.10 SE, *z* = 7.18, *p* < 0.001) and *pigeon* (β = 0.49 ± 0.09 SE, *z* = 5.31, *p* < 0.001) images, but not more than in *tortoise* images (β = 0.23 ± 0.10 SE, *z* = 2.31, *p* = 0.142), which in turn received more fixations than *deer* images (β = 0.47 ± 0.11 SE, *z* = 4.28, *p* < 0.001).

### Attention score

Overall, chimpanzees spent significantly more time looking at snakes than at carcasses (GLMM; β = 1.16 ± 0.07 SE, *z* = 2.12, *p* = 0.034; Fig. 3C, D; Table S9). In contrast, the proportion of time spent looking at the animals (versus elsewhere) was significantly lower in control images than in *carcass* images (*tortoise*: β = −0.25 ± 0.12 SE, *z* = –2.15, *p* = 0.032; *pigeon*: β = −0.57 ± 0.11 SE, *z* = –5.25, *p* < 0.001; *deer*: β = −0.79 ± 0.12 SE, *z* = –6.83, *p* < 0.001). Pairwise post hoc comparisons showed that both carcasses and snakes elicited longer total looking times than deer and pigeons (*carcass*–*deer*: β = 0.79 ± 0.12 SE, *z* = 6.83, *p* < 0.001; *carcass*–*pigeon*: β = 0.57 ± 0.11 SE, *z* = 5.25, *p* < 0.001; *snake*–*deer*: β = 0.94 ± 0.12 SE, *z* = 8.08, *p* < 0.001; *snake*–*pigeon*: β = 0.72 ± 0.11 SE, *z* = 6.55, *p* < 0.001). Snakes were also looked at for longer than tortoises (β = 0.40 ± 0.12 SE, *z* = 3.44, *p* = 0.005), which in turn received longer looking times than deer (β = 0.54 ± 0.13 SE, *z* = 4.10, *p* < 0.001). However, the difference in total looking time between carcasses and snakes was not significant in the post hoc tests (β = −0.16 ± 0.07 SE, *z* = –2.12, *p* = 0.210; Table S10), suggesting that the main effect detected by the GLMM was influenced by contrasts among several control conditions.

## DISCUSSION

Taken together, our results suggest that disgust-related visual and olfactory cues may interfere with cognitive performance in chimpanzees, whereas fear-relevant cues such as images of snakes appear to enhance performance and sustain attention. These findings support the hypothesis that fear of predators facilitates escape (or fight) by optimizing cognition and maintaining attention (*5, 6*), whereas disgust toward pathogen cues promotes avoidance through initial attention and cognitive disruption (*46, 59–61*).

Among the disgust-related images of Experiment 1, heterospecific animal carcasses selectively impaired performance in the number ordering task, reducing accuracy relative to disease-associated invertebrates. When considering task efficiency (/latency), chimpanzees completed trials faster under rotten food images, particularly at the beginning of a session. In the other conditions, their latency remained relatively stable over time, suggesting possible habituation or learning effects specific to the rotten food stimuli. Furthermore, chimpanzees may be less averse to spoiled food cues, particularly with repeated exposure. These differences may reflect variation in perceived threat or disease transmission routes. Animal carcasses, for instance, are among the strongest human disgust elicitors (*58, 62*), perhaps due to their association with death and salient sensory cues. Humans also rate death-related images as more disgusting than frightening (*58*). The absence of strong effects for invertebrate images may relate to distinct underlying defense systems – such as skin versus gut immunity – which trigger different behavioral responses (*63, 64*). While our statistical models accounted for subject variation, we noted striking behavioral responses in one female (Pendesa), who turned her back to the screen or knocked on the window when viewing disease-associated invertebrates (see Movie S4), suggesting an influence of subject experience. In humans, disgust is known to operate across functionally distinct domains (*65, 66*); whether such distinctions exist in chimpanzees remains an open and intriguing question.

Pathogen-associated odors, such as cadaverine and butyric acid, showed dynamic effects on task performance across sessions. During the first exposure, both odors were associated with shorter latencies to complete trials relative to water, suggesting a transient facilitation of performance that may reflect heightened arousal or an attempt to reduce exposure to the aversive stimulus. However, this effect did not persist: by the third session, butyric acid in particular was linked to significantly longer latencies, indicating reduced efficiency. Interestingly, a transient slowing also occurred early in the second session with limonene, a salient non-pathogen-related odor, suggesting that olfactory interference rather than aversion alone can momentarily affect performance. Taken together, these temporal shifts suggest that pathogen-related cues may initially elicit an alert or avoidance-oriented response, but later promote withdrawal or cognitive interference upon repeated exposure (*13,61,67*). While cadaverine could theoretically indicate both disease and predation risk, the overall pattern is more consistent with disgust than fear responses. In line with human findings showing that brief, high-concentration exposure to putrescine elicits faster reaction times (interpreted as heightened vigilance) (*67*), our lower concentrations and longer exposure revealed a later slowing and more cautious responding, reflecting the temporal dynamics of risk processing.

Both substances appear to elicit innate aversion in human neonates and fish (*68,69*), possibly due to the risks they signal: intoxication (butyric acid) and death (cadaverine). That cadaverine affected the behavior of captive chimpanzees – who may or may not have prior exposure to decaying flesh – supports the interpretation of an evolved, rather than learned, response. Cadaverine and putrescine are two of the main volatile organic compounds released during the decomposition of flesh. Prior studies have shown that these compounds are aversive to non-scavenging species, including humans (*67*), chimpanzees (*51*), and zebrafish (*69*), and can induce burial behavior in rats (*70*). In chimpanzees, this translates by spending less time near an object with putrescine compared to ammonia or water (*51*), although at much higher concentrations (3.5 g on cotton pads) than used here. Despite our lower concentration (0.3% for both cadaverine and butyric acid), we still observed some participation refusals (11% of experimental days), primarily for these two odors. We found no significant differences in performance between cadaverine and butyric acid, suggesting that they may be perceived similarly and belong to the same functional category of disgust-related cues.

Repeated exposure to carcass images significantly increased the latency to complete trials compared to snake images. These contrasting trends were not significant relative to their respective controls but reveal distinct temporal dynamics across emotional categories. Initially, chimpanzees responded faster under carcass images and slower under snake images compared to controls. However, after the fourth exposure (Fig. 2C), a divergence emerged: latencies for carcass images continued to increase, whereas latencies for snake images steadily decreased. This pattern suggests a differential effect of repeated exposure. The first exposure to snake images may have induced heightened alertness or caution, temporarily slowing responses. Over repeated exposures, this heightened state could have enhanced cognitive function – leading to faster reaction times, improved focus, or more effective decision-making – as chimpanzees became better at processing ecologically-relevant threats. An alternative explanation is habituation, but this seems less likely since different snake species and images were presented across trials and sessions, reducing stimulus familiarity. This differs from the improved performance seen with repeated exposure to rotten food images, where greater familiarity and prior experience may facilitate habituation or learning effects. The divergent patterns observed – faster responses to repeated fear-related stimuli (snakes) and slower responses to disgust-related stimuli (carcasses) – mirror findings in humans (*9,71,72*). For instance, Krusemark and Li (*46*) showed that fear-related images (snakes, spiders, but see (*73*), and guns) decrease reaction times, while disgust-related images (cockroaches, feces and vomit) increase them in a visual search task. This contrast supports the interpretation that fear can facilitate vigilance and rapid information processing, whereas disgust may induce avoidance or cognitive interference. Our data thus parallel human evidence and point to a shared evolutionary architecture for emotion-cognition interactions in chimpanzees. Furthermore, evidence of snake-sensitive neurons deep in the visual system of non-human primates (*74*) supports the idea of an evolved mechanism for rapid detection and efficient processing of threatening stimuli.

Snake and carcass images both captured visual attention more effectively than control images, as indicated by first fixation location and the number (proportion) of fixations – consistent with previous findings showing faster detection of threatening versus non-threatening stimuli in humans (*75–77*) and macaques (*78, 79*). On average, chimpanzees allocated about 71% of their viewing time to snakes and 67% to carcasses, compared to 62% for tortoises, 54% for pigeons and 49% for deer. This pattern indicates that both fear- and disgust-relevant cues strongly maintained visual attention. Although the difference between snakes and carcasses was not significant, snakes tended to elicit slightly more sustained attention, possibly because snakes represent an active threat that could approach and attack, whereas carcasses are inert. Indeed, features displayed in the carcass images (e.g. visible bones, smashed body, viscera) may have signaled to the chimpanzees that the carcass would not move. Previous eye-tracking studies have also shown that chimpanzees pay more attention (i.e. higher fixation durations) to death-related stimuli, such as conspecific skulls, compared to other species’ or control stimuli (*80*). Eye-tracking studies in humans show that disgusting images initially attract gaze (for about the first two seconds or the first trial) and subsequently trigger attentional avoidance (*12, 13, 61*), likely reflecting an evolved tendency to first assess threat and then avoid potential contamination. To clarify the temporal dynamics of chimpanzees’ attention to carcass images, future analyses should examine gaze allocation across time. In contrast, snake images are known to both capture and hold visual attention in human (*24,81*) and non-human primates (*82*), though see (*83*), which aligns with the hypothesis that snakes have been a key evolutionary threat for primates (*84, 85*).

Interestingly, we also observed significant differences across the control conditions, with tortoises receiving more fixations and being looked at longer than deer. Reptiles such as tortoises may attract more visual attention due to their distinctive morphology (e.g. the shell), which is evolutionarily more distant from primates and therefore may appear more novel or ambiguous with respect to potential threat. Elsewhere, Testudines have been used as control stimuli to test the snake detection theory using diverse methodologies such as immersive virtual reality, attentional interference tasks, and emotional/physiological ratings, with findings consistently showing that compared to Testudines, snakes elicit stronger avoidance behaviors, greater attentional capture, and more negative affective and physiological responses (*86–88*). However, other research suggests that features shared with snakes, such as reptilian scale patterns, may themselves trigger enhanced visual attention, raising the possibility that tortoises could partially activate evolved threat-detection mechanisms due to their morphological similarity (*89*).

### Limitations

To our knowledge, this study is the first to investigate cognitive processes under pathogen- and predator-related cues in a non-human primate. Our findings are therefore bounded in scope. For example, the fear-related effects reported here were elicited by snake images; it remains to be tested whether similar responses would emerge for other predator cues (*90*). Likewise, although we included pathogen-related odors, logistical constraints precluded the use of predator odors (e.g., felid urine; see (*91*)). Incorporating these cues in future work would extend the comparative framework across sensory modalities. Finally, despite a relatively large and diverse stimulus set (90 images) in the eye-tracking experiment, luminance was not standardized across images, which limited our ability to analyze pupil size.

### Conclusion and Perspectives

Our study provides novel evidence of how chimpanzees process pathogen- and predator-related cues in cognitive and eye-tracking tasks, revealing distinct patterns of facilitation and disruption in performance, alongside heightened attention compared to controls. These findings highlight the per-ceptual salience of ecologically relevant threats and point to promising avenues for future research. For instance, combining pathogen- and predator-related cues, whether within or across sensory modalities, could reveal how multiple threats jointly shape risk perception and behavior. Comple-mentary tools such as ChimpFACS (*92*) may help to identify specific facial muscle movements associated with exposure to carcass or snake stimuli. Further, comparing responses to live versus dead snakes would clarify whether animacy strengthens fear responses. Finally, examining how emotions spread socially – for example, whether observing conspecifics’ reactions influences an individual’s own avoidance or vigilance – will be essential to understanding the broader dynamics of threat perception.

## MATERIALS AND METHODS

### Subjects

We conducted four experiments with nine adult chimpanzees (*Pan troglodytes* spp.; 6 females and 3 males; 35.7 ± 12.7 y.o.; see SI Appendix Table S1) at the Center for the Evolutionary Origins of Human Behavior (EHUB; formerly named Primate Research Institute) of Kyoto University, Inuyama campus, Japan between October 2020 and January 2022, with an amendment in August 2023 to include session 3 of the carcass condition, which had not been conducted previously. Not all subjects participated in all four experiments, as inclusion depended on task suitability. For the number ordering task, chimpanzees were included only after achieving ≥ 90% accuracy in three consecutive training sessions. In addition, one participant (Pendesa) was excluded from the eye-tracking experiment because an arachnoid cyst in her brain appeared to have partially affected her visual field (see (*93*)).

### Ethics

The experimental protocol was approved by the Ethics Committee of the Primate Research Institute, Kyoto University (Approval #2019-212). All testing was conducted in dedicated indoor experimental booths (approx. 8 m³) located in the institute’s basement. Chimpanzees were called from their enriched outdoor enclosures and could freely choose whether to participate. If subjects stopped participating, the session was repeated when next possible. Food rewards were provided after each correct trial (Experiments 1–3), and juice was offered ad-libitum during eye-tracking sessions (Experiment 4). Each subject participated in brief sessions lasting 10–15 minutes, held in the morning or afternoon, and was not tested more than once per day under the same condition, but could be tested once in the morning and once in the afternoon under different conditions. The experimenters wore protective clothing and masks during tests to protect chimpanzees and themselves from potential exchange of pathogens. Experimental booths were cleaned after each subject experimental session.

### Apparatuses

Experiments 1–3 involved a number ordering task that chimpanzees at EHUB have performed for more than two decades (*52,94–96*). Nine numerals (one to nine) except for one subject (Pendesa: one to six) were presented at random locations on a 17” LCD touch screen (1280 × 1024 px) during one given trial. Chimpanzees had to touch the numerals in an ascending order. If correct, they received a small piece of apple or a dry raisin from a feeder connected to the computer running the task via Visual Basic 2010 (Microsoft Corporation, Redmond, Washington, USA). These experiments were video-recorded from outside the booth by two GoPro Hero 8 Black cameras attached above the screen and on the side of the booth. Experiment 4 consisted of a visual fixation task using an eye-tracker (300 Hz; X300; Tobii Technology AB, Stockholm, Sweden) connected to a laptop (Galleria GR2060RGF-T) with its software (Tobii Pro Lab v.1.181.37603), a method originally developed with humans, now widespread with other great ape species (*97, 98*). Chimpanzees were 60-70 cm away from a screen (16:9; 1920 × 1080 px) placed behind the Plexiglas of the booth, with their head held straight using a grape juice-sucking system (described in Kano & Call (*99*)) involving a drip bag and tube inserted into a hole in the Plexiglas of the booth. The light of the room was turned off and the subject partially isolated from visual distractors by covering the side walls of the booth with black curtains (Fig. S5).

### Experiment 1: Disgust-related images

We tested the influence of disgust-related images (DIRTI) on six chimpanzees’ performance in the number ordering task. To do this, we selected free right DIRTI from Haberkamp et al. (*58*) (https://zenodo.org/records/167037), and from free (Unplash) and paid (iStock, Adobe Stock) image platforms. We selected three conditions of disgust elicitors based on disgust ratings in humans (*58*) and their relevance to chimpanzees: non-conspecific animal carcasses (from mammals, birds, amphibians, and marsupials, showing typical signs of decay such as tissue breakdown, presence of maggots, visible skeleton), disease-associated (e.g. mosquito, leech) or -mimicking (e.g. earthworm, slug) invertebrates, and rotten food (spoiled fruits and vegetables) (Fig. S1-S3). Control image conditions were sleeping animals, invertebrates not associated with disease risk and edible food items, respectively. One session consisted of 59 trials of the number ordering task. It started with nine warm-up trials, followed by the full-screen presentation of a disgust-related or control image (1260 × 945 mm) for one second. This image was then followed by one test trial and four baseline trials, resulting in 10 image presentations per session. Each chimpanzee completed one randomized image session per condition per day. Each condition was repeated across three sessions per subject, and each session consisted of a different set of images (i.e. 3 sets of 10 different images per condition). In total, chimpanzees performed 108 sessions (3 sessions × 6 image conditions × 6 subjects).

### Experiment 2: Pathogen-associated odors

Here, we tested whether pathogen-associated odors (PATOS) had a distracting effect on chimpanzee cognitive performance, using a paradigm similar to that used for DIRTI. Specifically, we used two volatile organic compounds previously employed in aversion research with humans and zebra fish (*69, 100–102*): cadaverine, associated with body decomposition; and butyric acid, associated with vomit and illness. Two control odors, limonene and water, were also included. Odors were delivered via a custom-built olfactory phenotyping device (“Smox”) placed beneath the touch screen and behind a thin grid. The device consisted of a 27 × 26 × 15 cm polycarbonate box containing a nebulizer (Omron NE-U100) with 6 mL of water or odor solution: 6 mL of water + 150 µL limonene (2.5% concentration) / 20 µL cadaverine (1,5-Diaminopentane) / butyric acid (both at 0.3% concentration) (all Tokyo Chemical Industry), and a small battery-powered fan on top (Fig. 2B). Odor concentrations were chosen based on pilot tests with human experimenters to ensure they were detectable yet tolerable. Each session consisted of 50 trials of the number ordering task under continuous odor diffusion, starting upon the chimpanzee entering the booth; therefore no warm-up trials were included. To maintain task engagement, we increased the reward’s nutritive value (i.e. piece of apple plus berries) and reduced the number of numerals shown (from 9 to 6). A maximum of two chimpanzees were tested per PATOS per day, with at least a four-hour interval (except for water) and the use of an air purifier (LEAF 320i) between sessions. Each chimpanzee completed three sessions per odor condition (72 sessions in total).

### Experiment 3: Carcass versus snake images

We applied the same design as in Experiment 1, using four image conditions: non-conspecific animal carcasses (new set of images), snakes, and scrambled mosaics of both image types (Fig. 1C). The scrambled mosaics were generated using WebMorph (*103*) by dividing the original images into 12 × 9 squares. These mosaics preserve low-level visual features while disrupting the global structure of the image, a manipulation shown to significantly reduce the activation of pulvinar neurons in response to snake stimuli (*74*). The same six chimpanzees as in Experiment 1 participated (Table S1), completing a total of 72 sessions (3 sessions × 4 image conditions × 6 subjects).

### Experiment 4: Eye-tracking

We presented a total of 90 images to 8 chimpanzees ([10 images x 2 test categories x 3 sessions] + [10 images x 3 control categories x 1 session]). The test set included non-conspecific animal carcasses and snakes (new sets of images), while the control sets featured live ungulates (deer), birds (pigeons) and tortoises, which are not known to be associated to specific threats to chimpanzees. Each image featured one single animal, full body from front, top or profile in a natural background environment. The images were positioned centrally within a grey horizontal slide, occupying 25% of the screen. Each session started with eye calibration, followed by the display of the first image (of one category) for five seconds. Following this, three fixation points, located in one corner of each slide (such as one cross, one number and one dot in the top right corner) would be presented one after another by switching via key press. Subsequently, the second image of the same category would be displayed for five seconds, followed by the relocation of the three fixation points to another corner of the slide, and this pattern would continue until the presentation of the tenth image, ending the session (Movie S5). The same image order for each set was presented to the chimpanzees. For each picture slide, we defined three areas of interest (AOIs): 1) the animal, 2) the environment surrounding it, and 3) the grey area surrounding the picture frame (Fig. S4). We considered three proxies of visual attention capture and retention, obtained via Tobii Pro Lab: 1) first fixation location, 2) proportion of fixations, and 3) attention score (i.e. the proportion of time looking at the animal versus elsewhere). We redid a session if it contained below 30% of fixations or if the subject left the experimental room, both happening in less than 10% of all sessions. A few exceptions were made: Pal’s fourth attempt at the “carcass” third session, which resulted in only 25% of fixations; Akira’s “snake” first and third sessions, where he left midway after several failed attempts; and Akira’s “deer” session, for which he never passed calibration. We therefore retained Pal’s final attempt at the “carcass” third session but excluded Akira’s “snake” first and third, and “deer” sessions. The low threshold (percentage of fixations) retained is voluntary and due to the prediction, associated with the task, that DIRTI elicit gaze avoidance (*13*). After data filtering based on these conditions, we obtained 47 recordings for the test sessions and 23 recordings for the control sessions.

### Data analyses

We analyzed how disgust- and fear-related sensory cues influenced chimpanzee cognitive processes using generalized linear mixed models (GLMMs), implemented via the *glmmTMB* package (*104*). In Experiments 1–3, response variables were trial accuracy (binary; correct = 1, incorrect = 0), modeled with a binomial distribution; reaction time (continuous; latency to touch the first numeral); and latency to complete correct trials (continuous), both modeled with Gamma distributions. Only data from after the initial nine warm-up trials were included in analyses for Experiments 1 and 3. We excluded trials with latencies below 300 ms (indicative of anticipatory responses or hands already on the screen) or above 2500 ms (suggesting distraction or disengagement). Control conditions in Experiment 1 (images of animals sleeping, non-pathogenic invertebrates and edible food items) were tested for equivalence in success rates via a Chi-squared test before being grouped into a single control category (see Results). Predictor variables included condition (i.e. the different images or odors), trial number (1-50; scaled and included in the accuracy models for Experiments 1–3 and the reactivity and latency models for Experiment 2), number of image exposure (1-10; scaled and included in the reactivity and latency models for Experiments 1 and 3), and session number (1-3; included in the reactivity and latency models for Experiment 2). Note that for the reactivity and latency models of Experiments 1 and 3, only the trials immediately following image exposure were retained for data analysis. While the accuracy models account for potential longer-lasting effects of the images on cognition, the reactivity and latency models are designed to capture the immediate effect of image exposure. All models included a random intercept for subject identity to account for repeated measures on the same participants. In addition, models included a nested random effect of subjects within testing dates to account for potential variability across sessions; this nested effect was omitted from the accuracy model in Experiment 1 because it complicated model fitting without improving fit.

In Experiment 4, the response variables included the location of the first fixation (binary; animal vs. other); the proportion of fixations on the animal (continuous; bounded between 0 and 1) and the proportion of time spent looking at the animal (continuous; bounded between 0 and 1). Proportion data were modeled using beta distributions. There was one predictor variable: condition (image category) and three random effects: the slide number per session to account for variability in the image presented, subject identity and observation (row ID; for proportion models only).

Model comparisons were conducted using likelihood ratio tests (LRTs) from the *lmtest* package (*105*), comparing full models (including all predictors) against null models with only control variables. We also tested models with and without interactions of interest (e.g. condition and trial number). Model diagnostics were performed using the *DHARMa* package (*106*) to check residual homogeneity, homoscedasticity, and zero-inflation; no violations of model assumptions were found, except for the *Proportion of fixations* model, which indicated a slight deviation from uniform residual distribution (Kolmogorov-Smirnov test: D = 0.055, *p* = 0.030). Despite this, the model provided the best fit among several alternatives (lower AIC, more stable estimates), and residual patterns did not suggest substantial model misspecification. We therefore retained this model for inference, while acknowledging minor limitations in model fit. For all retained models, we conducted pairwise post hoc comparisons among conditions using the *emmeans* package (*107*), applying Tukey correction (or Bonferroni correction for pre-selected contrasts in Experiment 3). All analyses were conducted in R v.4.4.2 (*108*).

## Supplementary Materials

Tables S1 to S10

Figs. S1 to S5

Movies S1 to S5

## Acknowledgments

We thank the chimpanzee caretakers Akemi Hirakuri, Etsuko Ichino and Tomoko Takashima for their help with the experiments. C.S. would like to thank Yena Kim, Kenneth Keuk and Sylvain Dorrière for insightful discussions regarding data analyses and eye-tracking. C.S. would also like to thank Frans de Waal for stopping by her poster at IPS 2018 in Chicago and inspiring the exploration of the cognitive facet of disgust in non-human primates. **Funding.** This work was supported by the Japan Society for the Promotion of Science (Postdoctoral Fellowship P19088 to C.S.). C.S. also acknowledges funding from the Institute for Advanced Study in Toulouse (IAST) through the French National Research Agency (ANR) under grant ANR-17-EURE-0010 (Programme d’Investissements d’Avenir), and from a Marie Sklodowska-Curie Postdoctoral Fel-lowship funded by UK Research and Innovation (UKRI) under the UK government’s Horizon Europe funding guarantee, grant number EP/Z001625/1. **Author contributions.** C.S. designed research and analysed data. C.S., A.G. and I.A. realised experiments. I.A., A.J.J.M. and N.K. supervised research. C.S. wrote the paper and N.K., A.J.J.M., A.G., and I.A. reviewed and edited it. **Competing interests.** The authors declare no conflicts of interest. **Data availability.** All study data are available in Github (https://github.com/CSarabian/chimp-dead-vs-death.git) and Supplementary Materials.

