## Supplementary material for "Dissecting disgust and fear in chimpanzees: From facilitation to disruption of cognitive processes": Sarabian_et_al_SM.pdf

**Table S1**: **Participating subject details.** Subjects were nine chimpanzees from Kyoto University Primate Research Institute, involved in Experiments 1–4. Sex is given as m for males and f for females, and age is reported in years at the time of the study.

| Subject | Sex | Age | Experiment |
|---|---|---|---|
| Ai | F | 44 | 1–4 |
| Akira | M | 44 | 4 |
| Ayumu | M | 20 | 1–4 |
| Chloe | F | 40 | 1–4 |
| Cleo | F | 20 | 1–4 |
| Gon | M | 54 | 4 |
| Pal | F | 20 | 1–4 |
| Pan | F | 36 | 4 |
| Pendesa | F | 43 | 1–3 |

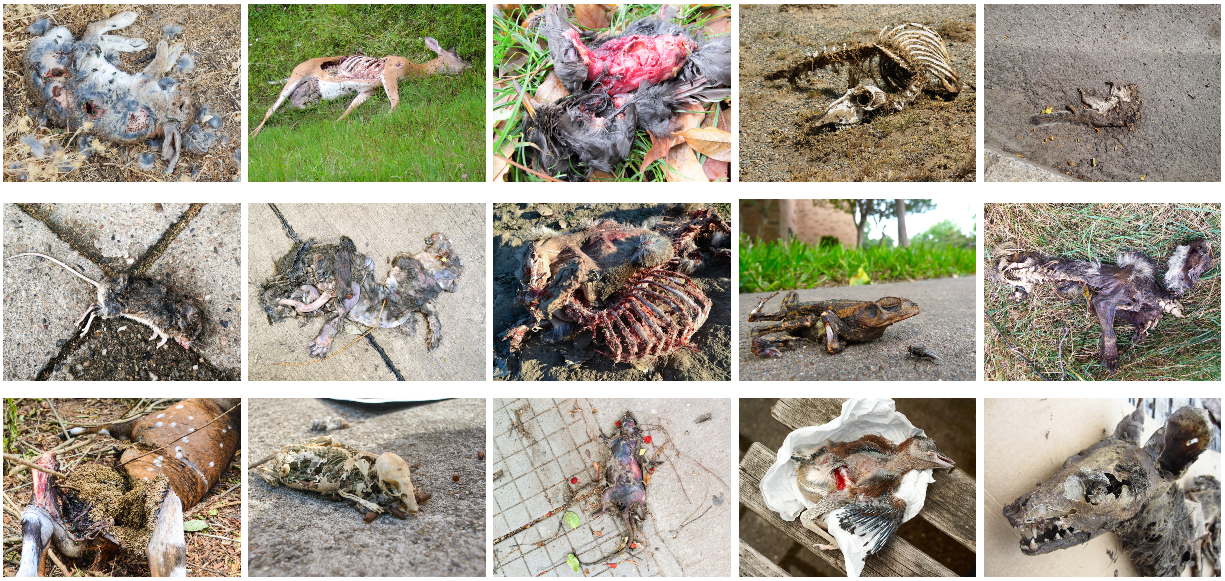

**Fig. S1**: **Animal carcass images used in Experiment 1.** Each row displays images from the same session. Those shown here are from the DIRTI database (https://zenodo.org/records/167037). Five additional images per session came from paid platforms and are not available for redistribution.

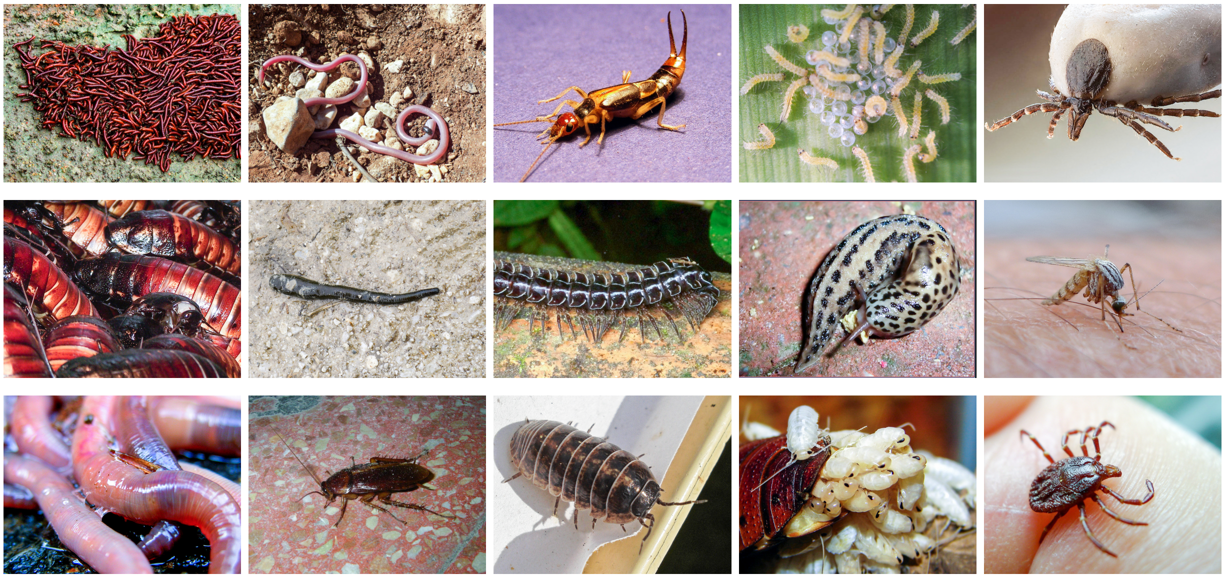

**Fig. S2**: **Disease-associated/pathogen-mimicking invertebrate images used in Experiment 1.** Each row displays images from the same session. Those shown here are from the DIRTI database (https://zenodo.org/records/167037). Five additional images per session came from paid platforms and are not available for redistribution.

**Table S2**: **Likelihood ratio tests comparing full models vs null and/or additive models for Experiments 1-4 (M1-4) for the different response variables (MX.1-3).** Full models include predictors of interest or interactions between predictors. Null models contain only control predictors or the intercept. Additive models contain predictors of interest without interaction terms. Bold text denotes models that significantly outperformed their respective null or additive models. Significance levels: ***$p$ < 0.001, **$p$ < 0.01, *$p$ < 0.05.

| Statistical Model [X] | ΔLogLik | Δd.f. | $X^2$ | $p$-value |
|---|---|---|---|---|
| **Experiment 1 (DIRTI)** | | | | |
| **[M1.1] Accuracy** | | | | |
| **Condition*Trial vs Trial** | **8.10** | **6** | **16.24** | **0.013*** |
| Condition*Trial vs Condition + Trial | 3.60 | 3 | 7.34 | 0.062 |
| **Condition + Trial vs Trial** | **4.50** | **3** | **8.90** | **0.031*** |
| [M1.2] Reactivity | | | | |
| Condition*Exposure vs Exposure | 5.10 | 6 | 10.27 | 0.114 |
| **[M1.3] Latency** | | | | |
| **Condition*Exposure vs Exposure** | **9.40** | **6** | **18.87** | **0.004**** |
| **Condition*Exposure vs Condition + Exposure** | **7.00** | **3** | **13.97** | **0.003**** |
| **Experiment 2 (PATOS)** | | | | |
| [M2.1] Accuracy | | | | |
| Odor*Trial*Session vs Trial + Session | 6.10 | 13 | 12.25 | 0.507 |
| [M2.2] Reactivity | | | | |
| Odor*Trial*Session vs Trial + Session | 9.00 | 13 | 17.12 | 0.194 |
| **[M2.3] Latency** | | | | |
| **Odor*Trial*Session vs Trial + Session** | **16.00** | **13** | **33.64** | **0.001**** |
| **Odor*Trial*Session vs Odor + Trial + Session** | **15.00** | **10** | **31.37** | **< 0.001***** |
| **Experiment 3 (Carcass vs snake images)** | | | | |
| [M3.1] Accuracy | | | | |
| Condition*Image*Trial vs Trial | 5.80 | 6 | 11.53 | 0.073 |
| [M3.2] Reactivity | | | | |
| Condition*Image*Exposure vs Exposure | 4.10 | 6 | 8.21 | 0.223 |
| **[M3.3] Latency** | | | | |
| **Condition*Image*Exposure vs Exposure** | **7.10** | **6** | **14.27** | **0.027*** |
| **Condition*Image*Exposure vs Cond + Image + Expo** | **6.90** | **4** | **13.77** | **0.008**** |
| **Experiment 4 (Eye-tracking)** | | | | |
| **[M4.1] First fixation location** | | | | |
| **Condition vs intercept-only model** | **14.92** | **4** | **29.84** | **< 0.001***** |
| **[M4.2] Proportion of fixations** | | | | |
| **Condition vs intercept-only model** | **32.68** | **4** | **65.37** | **< 0.001***** |
| **[M4.3] Proportion of time looking** | | | | |
| **Condition vs intercept-only model** | **42.68** | **4** | **85.36** | **< 0.001***** |

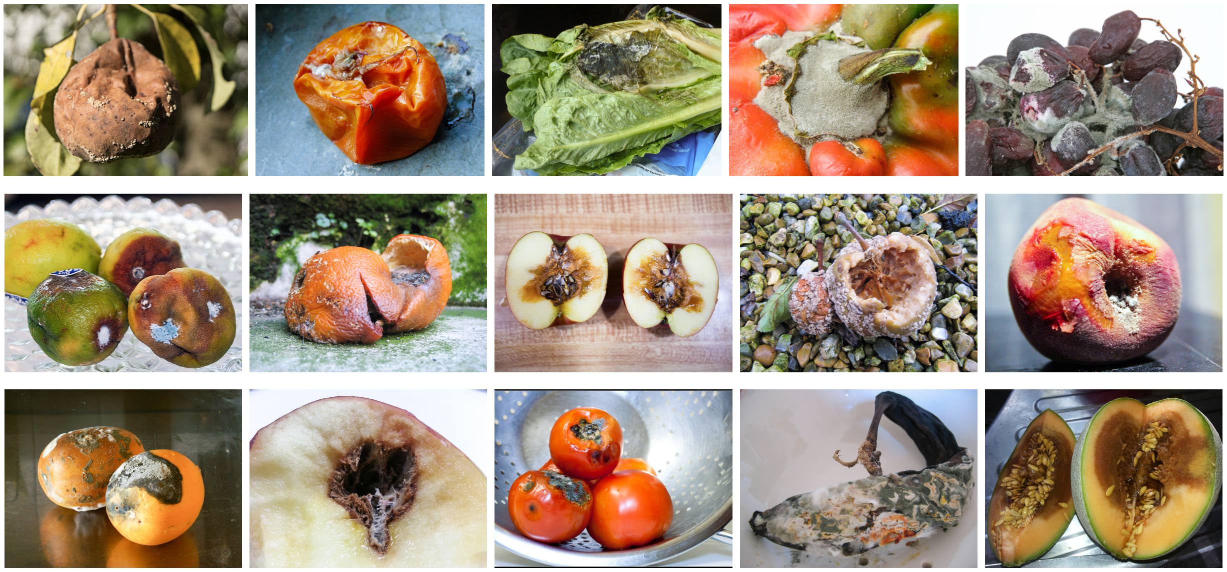

**Fig. S3**: **Rotten food images used in Experiment 1.** Each row displays images from the same session. Those shown here are from the DIRTI database (https://zenodo.org/records/167037). Five additional images per session came from paid platforms and are not available for redistribution.

**Table S3**: **Summary of GLMM results for predictors of interest in Experiment 1.** The reference level is the pooled control image condition. Significant effects are highlighted in bold. Significance levels: ***$p < 0.001$, **$p < 0.01$, *$p < 0.05$.

| Model | Predictor | Estimate ($\beta$) | SE | $z$ | $p$-value |
|---|---|---|---|---|---|
| [M1.1] Accuracy | (Intercept) | 1.691 | 0.166 | 10.17 | $< 2e - 16 * **$ |
| | **Carcass** | **-0.244** | **0.100** | **-2.44** | **0.015*** |
| | Invertebrate | 0.105 | 0.109 | 0.96 | 0.335 |
| | Rotten food | -0.096 | 0.103 | -0.93 | 0.354 |
| | Trial number | 0.037 | 0.037 | 0.99 | 0.323 |
| [M1.3] Latency | (Intercept) | 8.312 | 0.051 | 164.43 | $< 2e - 16 * **$ |
| | Carcass | -0.008 | 0.019 | -0.44 | 0.657 |
| | Invertebrate | -0.021 | 0.018 | -1.17 | 0.241 |
| | Rotten food | 0.030 | 0.018 | 1.64 | 0.102 |
| | Exposure | -0.002 | 0.006 | -0.29 | 0.770 |
| | Carcass*Exposure | 0.015 | 0.013 | 1.16 | 0.247 |
| | Invertebrate*Exposure | 0.003 | 0.013 | 0.23 | 0.816 |
| | **Rotten food*Exposure** | **-0.041** | **0.013** | **-3.13** | **0.002**** |

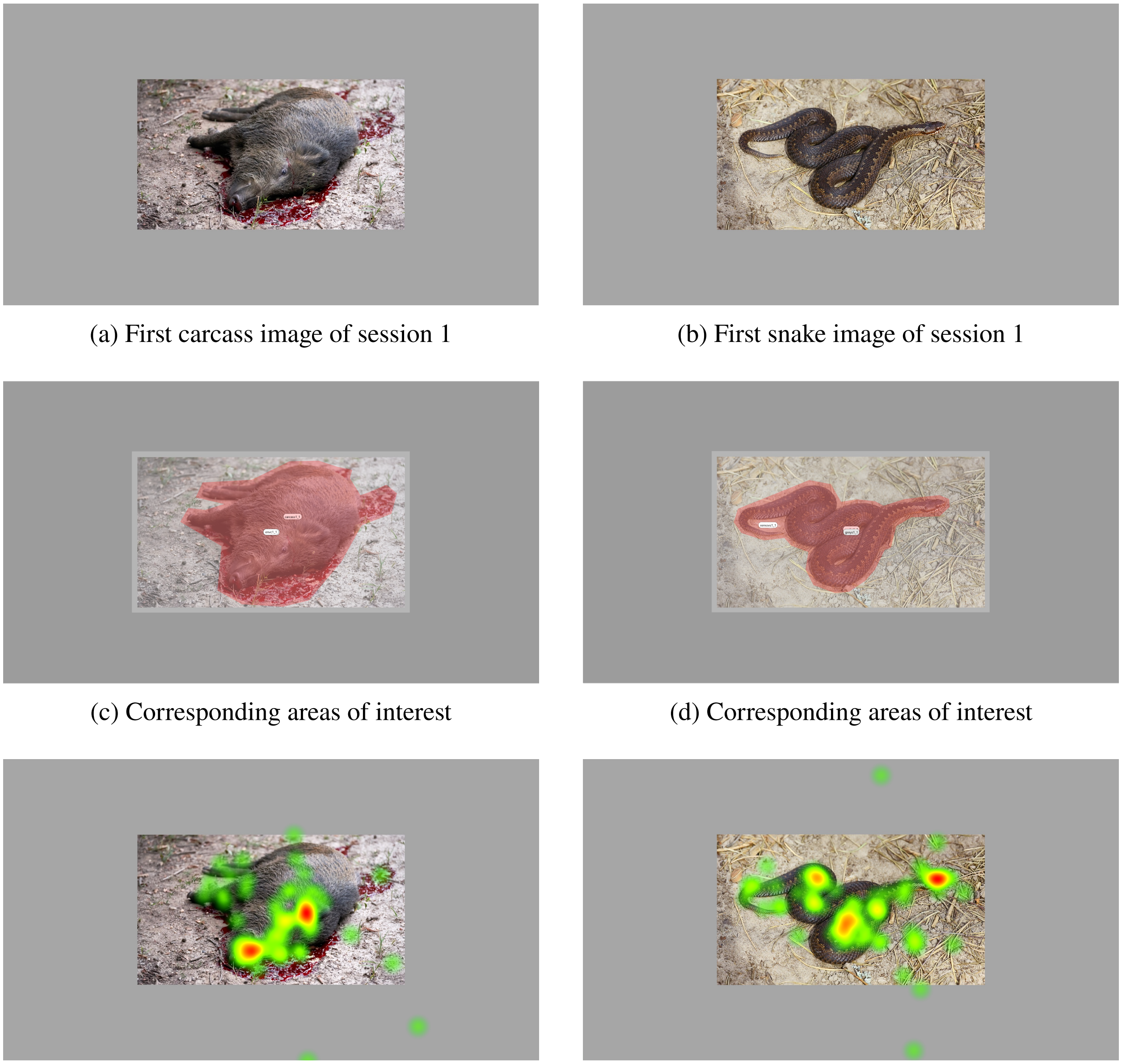

(a) First carcass image of session 1

(b) First snake image of session 1

(c) Corresponding areas of interest

(d) Corresponding areas of interest

(e) Corresponding heat map

(f) Corresponding heat map

**Fig. S4**: **Areas of interest considered in Experiment 4.** These included the animal, its environment and the grey area surrounding the picture frame, as illustrated in (c) and (d) for images (a) and (b), respectively. The heat maps (e) and (f) represent the corresponding fixation counts for all chimpanzees; the “hotter” areas are where participants fixated more often.

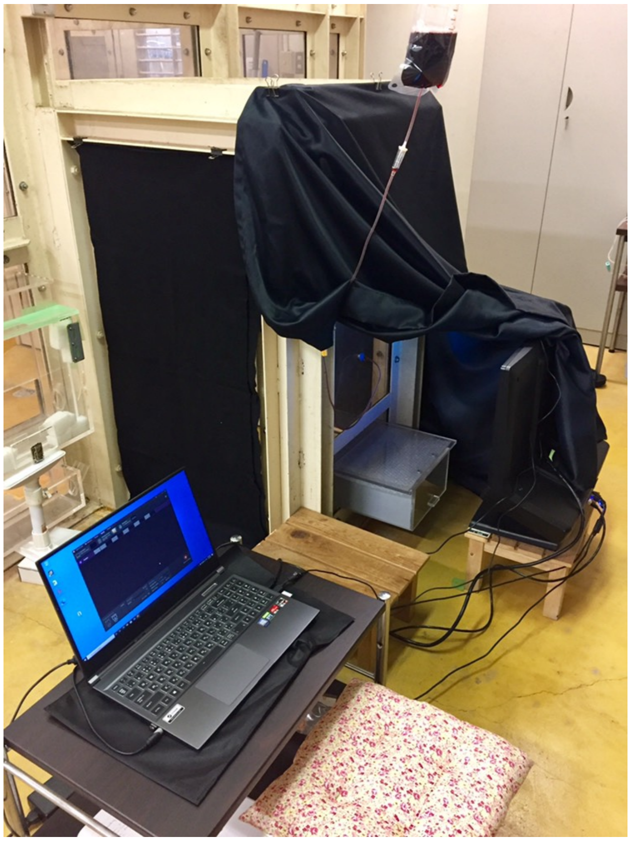

**Fig. S5**: **Experimental setup for Experiment 4.** Chimpanzees were positioned 60–70 cm from the monitor screen with their head stabilized using a grape juice–sucking system (drip bag and tube inserted through the Plexiglas panel). The testing room was darkened, and side walls of the booth were covered with black curtains to reduce visual distractions.

**Table S4**: **Estimated marginal means and pairwise comparisons of accuracy and latency across image conditions in Experiment 1.** For latency, analyses were performed at min, mid, and high image exposure. Image codes: Ctrl = Control; C = Carcass; I = Invertebrate; RF = Rotten food. Pairwise comparisons are Tukey-adjusted. Results for accuracy are shown on the logit scale, and those for latency on the log scale. Significant effects are highlighted in bold. Significance levels: $^{***}p < 0.001$, $^{**}p < 0.01$, $^{*}p < 0.05$.

| Model | Expo | Cond | EMM (95% CI) | Contrast | $\beta$ | SE | $z$ | $p$-value |
|---|---|---|---|---|---|---|---|---|
| [M1.1] Accuracy | | Ctrl | 1.69 [1.37, 2.02] | - | - | - | - | - |
| | | C | 1.45 [1.10, 1.80] | Ctrl - C | 0.244 | 0.100 | 2.44 | 0.070 |
| | | I | 1.80 [1.44, 2.16] | Ctrl - I | -0.105 | 0.109 | -0.96 | 0.770 |
| | | RF | 1.60 [1.24, 1.95] | Ctrl - RF | 0.096 | 0.103 | 0.93 | 0.791 |
| | | | | **C - I** | **-0.349** | **0.127** | **-2.74** | **0.031*** |
| | | | | C - RF | -0.149 | 0.123 | -1.21 | 0.622 |
| | | | | I - RF | 0.200 | 0.130 | 1.54 | 0.412 |
| [M1.3] Latency | Min | Ctrl | 8.31 [8.21, 8.42] | - | - | - | - | - |
| | | C | 8.28 [8.17, 8.39] | Ctrl - C | 0.033 | 0.028 | 1.18 | 0.641 |
| | | I | 8.29 [8.18, 8.40] | Ctrl - I | 0.026 | 0.027 | 0.96 | 0.773 |
| | | RF | 8.41 [8.30, 8.52] | **Ctrl - RF** | **-0.096** | **0.029** | **-3.34** | **0.005**** |
| | | | | C - I | -0.006 | 0.034 | -0.19 | 0.998 |
| | | | | **C - RF** | **-0.128** | **0.035** | **-3.69** | **0.001**** |
| | | | | **I - RF** | **-0.122** | **0.035** | **-3.53** | **0.002**** |
| | Mid | Ctrl | 8.31 [8.21, 8.41] | - | - | - | - | - |
| | | C | 8.30 [8.20, 8.41] | Ctrl - C | 0.008 | 0.019 | 0.44 | 0.971 |
| | | I | 8.29 [8.19, 8.39] | Ctrl - I | 0.021 | 0.018 | 1.17 | 0.644 |
| | | RF | 8.34 [8.24, 8.44] | Ctrl - RF | -0.030 | 0.019 | -1.64 | 0.359 |
| | | | | C - I | 0.013 | 0.023 | 0.58 | 0.939 |
| | | | | C - RF | -0.039 | 0.023 | -1.69 | 0.331 |
| | | | | I - RF | -0.052 | 0.022 | -2.30 | 0.098 |
| | High | Ctrl | 8.31 [8.21, 8.41] | - | - | - | - | - |
| | | C | 8.32 [8.22, 8.43] | Ctrl - C | -0.016 | 0.028 | -0.55 | 0.947 |
| | | I | 8.29 [8.18, 8.40] | Ctrl - I | 0.017 | 0.027 | 0.61 | 0.929 |
| | | RF | 8.27 [8.17, 8.38] | Ctrl - RF | 0.034 | 0.027 | 1.27 | 0.584 |
| | | | | C - I | 0.032 | 0.034 | 0.94 | 0.783 |
| | | | | C - RF | 0.050 | 0.034 | 1.46 | 0.460 |
| | | | | I - RF | 0.017 | 0.033 | 0.53 | 0.953 |

**Table S5**: **Summary of GLMM results for predictors of interest in Experiment 2.** The reference odor is water. Significant effects are highlighted in bold. Significance levels: ***$p < 0.001$, **$p < 0.01$, *$p < 0.05$.

| Model | Predictor | Estimate ($\beta$) | SE | $z$ | $p$-value |
|---|---|---|---|---|---|
| [M2.3] Latency | (Intercept) | 7.946 | 0.047 | 168.10 | $< 2e - 16 ***$ |
| | Butyric acid | -0.095 | 0.054 | -1.77 | 0.077 |
| | Cadaverine | -0.054 | 0.051 | -1.04 | 0.297 |
| | Limonene | -0.036 | 0.051 | -0.70 | 0.487 |
| | **Trial** | **0.073** | **0.022** | **3.29** | **< 0.001*** ** |
| | Session | 0.031 | 0.017 | 1.85 | 0.064 |
| | **Butyric acid*Trial** | **-0.126** | **0.033** | **-3.78** | **< 0.001*** ** |
| | **Cadaverine*Trial** | **-0.119** | **0.033** | **-3.65** | **< 0.001*** ** |
| | Limonene*Trial | -0.060 | 0.032 | -1.89 | 0.059 |
| | **Butyric acid*Session** | **0.068** | **0.025** | **2.75** | **0.006*** * |
| | Cadaverine*Session | 0.030 | 0.024 | 1.25 | 0.211 |
| | Limonene*Session | 0.027 | 0.024 | 1.11 | 0.266 |
| | **Trial*Session** | **-0.041** | **0.010** | **-3.94** | **< 0.001*** ** |
| | **Butyric acid*Trial*Session** | **0.048** | **0.015** | **3.12** | **< 0.001*** ** |
| | **Cadaverine*Trial*Session** | **0.056** | **0.015** | **3.60** | **< 0.001*** ** |
| | Limonene*Trial*Session | 0.021 | 0.015 | 1.40 | 0.160 |

**Table S6**: **Estimated marginal means and pairwise comparisons from the GLMM testing the effects of odor, trial number, and session on latency in Experiment 2.** Analyses were performed at early, mid, and high trial numbers. Odor codes: W = Water; BA = Butyric acid; C = Cadaverine; L = Limonene. Pairwise comparisons are Tukey-adjusted. Significant effects are highlighted in bold. Significance levels: $^{***}p < 0.001$, $^{**}p < 0.01$, $^{*}p < 0.05$.

| Session – Trial | Condition | EMM (95% CI) | Contrast | $\beta$ | SE | $z$ | $p$-value |
|---|---|---|---|---|---|---|---|
| 1 – Early | W | 7.93 [7.84, 8.02] | - | - | - | - | - |
| | BA | 8.01 [7.92, 8.11] | W - BA | -0.083 | 0.051 | -1.63 | 0.364 |
| | C | 8.02 [7.92, 8.11] | W - C | -0.085 | 0.049 | -1.72 | 0.315 |
| | L | 7.95 [7.86, 8.04] | W - L | -0.020 | 0.048 | -0.41 | 0.977 |
| | | | BA - C | -0.002 | 0.052 | -0.04 | 1.000 |
| | | | BA - L | 0.063 | 0.051 | 1.25 | 0.598 |
| | | | C - L | 0.065 | 0.049 | 1.33 | 0.547 |
| 1 – Mid | W | 7.98 [7.91, 8.06] | - | - | - | - | - |
| | BA | 7.96 [7.88, 8.04] | W - BA | 0.029 | 0.035 | 0.82 | 0.845 |
| | C | 7.96 [7.89, 8.04] | W - C | 0.020 | 0.033 | 0.62 | 0.926 |
| | L | 7.97 [7.90, 8.05] | W - L | 0.014 | 0.033 | 0.44 | 0.972 |
| | | | BA - C | -0.008 | 0.035 | -0.23 | 0.996 |
| | | | BA - L | -0.014 | 0.035 | -0.41 | 0.977 |
| | | | C - L | -0.006 | 0.033 | -0.18 | 0.998 |
| 1 – High | W | 8.04 [7.95, 8.13] | - | - | - | - | - |
| | BA | 7.90 [7.81, 8.00] | **W - BA** | **0.138** | **0.050** | **2.75** | **0.031*** |
| | C | 7.91 [7.82, 8.01] | **W - C** | **0.124** | **0.048** | **2.57** | **< 0.05*** |
| | L | 7.99 [7.90, 8.08] | W - L | 0.048 | 0.047 | 1.02 | 0.740 |
| | | | BA - C | -0.014 | 0.051 | -0.28 | 0.993 |
| | | | BA - L | -0.090 | 0.050 | -1.80 | 0.273 |
| | | | C - L | -0.076 | 0.048 | -1.58 | 0.388 |
| 2 – Early | W | 8.00 [7.91, 8.09] | - | - | - | - | - |
| | BA | 8.14 [8.04, 8.24] | W - BA | -0.140 | 0.055 | -2.56 | 0.051 |
| | C | 8.03 [7.93, 8.12] | W - C | -0.024 | 0.051 | -0.48 | 0.964 |
| | L | 8.14 [8.05, 8.23] | **W - L** | **-0.139** | **0.051** | **-2.70** | **0.035*** |
| | | | BA - C | 0.116 | 0.055 | 2.10 | 0.153 |
| | | | BA - L | 0.001 | 0.056 | 0.03 | 1.000 |
| | | | C - L | -0.115 | 0.052 | -2.21 | 0.122 |
| 2 – Mid | W | 7.99 [7.91, 8.06] | - | - | - | - | - |
| | BA | 8.04 [7.95, 8.12] | W - BA | -0.047 | 0.038 | -1.25 | 0.596 |
| | C | 8.00 [7.92, 8.07] | W - C | -0.006 | 0.035 | -0.18 | 0.998 |
| | L | 8.01 [7.94, 8.09] | W - L | -0.025 | 0.035 | -0.70 | 0.898 |
| | | | BA - C | 0.041 | 0.039 | 1.06 | 0.716 |
| | | | BA - L | 0.023 | 0.039 | 0.58 | 0.938 |
| | | | C - L | -0.018 | 0.036 | -0.51 | 0.957 |

| Session – Trial | Condition | EMM (95% CI) | Contrast | $\beta$ | SE | $z$ | $p$-value |
|---|---|---|---|---|---|---|---|
| 2 – High | W | 7.98 [7.89, 8.07] | - | - | - | - | - |
| | BA | 7.93 [7.83, 8.04] | W - BA | 0.045 | 0.056 | 0.80 | 0.854 |
| | C | 7.97 [7.87, 8.06] | W - C | 0.012 | 0.050 | 0.23 | 0.996 |
| | L | 7.89 [7.79, 7.99] | W - L | 0.088 | 0.051 | 1.71 | 0.318 |
| | | | BA - C | -0.033 | 0.057 | -0.58 | 0.939 |
| | | | BA - L | 0.043 | 0.058 | 0.75 | 0.876 |
| | | | C - L | 0.076 | 0.053 | 1.45 | 0.471 |
| 3 – Early | W | 8.13 [8.04, 8.22] | - | - | - | - | - |
| | BA | 8.18 [8.09, 8.28] | W - BA | -0.053 | 0.051 | -1.03 | 0.733 |
| | C | 8.09 [7.99, 8.18] | W - C | 0.045 | 0.052 | 0.87 | 0.823 |
| | L | 8.11 [8.01, 8.21] | W - L | 0.018 | 0.054 | 0.33 | 0.987 |
| | | | BA - C | 0.097 | 0.054 | 1.80 | 0.274 |
| | | | BA - L | 0.071 | 0.056 | 1.26 | 0.589 |
| | | | C - L | -0.027 | 0.057 | -0.47 | 0.965 |
| 3 – Mid | W | 8.05 [7.97, 8.12] | - | - | - | - | - |
| | BA | 8.15 [8.07, 8.23] | **W - BA** | **-0.107** | **0.035** | **-3.04** | **0.013*** |
| | C | 8.08 [8.01, 8.16] | W - C | -0.038 | 0.035 | -1.07 | 0.708 |
| | L | 8.08 [8.00, 8.16] | W - L | -0.037 | 0.035 | -1.03 | 0.731 |
| | | | BA - C | 0.069 | 0.037 | 1.88 | 0.238 |
| | | | BA - L | 0.070 | 0.037 | 1.91 | 0.224 |
| | | | C - L | 0.001 | 0.037 | 0.04 | 1.000 |
| 3 – High | W | 7.96 [7.87, 8.05] | - | - | - | - | - |
| | BA | 8.12 [8.03, 8.22] | **W - BA** | **-0.160** | **0.051** | **-3.12** | **< 0.01**** |
| | C | 8.08 [7.98, 8.18] | W - C | -0.119 | 0.052 | -2.29 | 0.102 |
| | L | 8.05 [7.96, 8.15] | W - L | -0.090 | 0.051 | -1.76 | 0.294 |
| | | | BA - C | 0.041 | 0.054 | 0.76 | 0.875 |
| | | | BA - L | 0.070 | 0.054 | 1.31 | 0.558 |
| | | | C - L | 0.029 | 0.054 | 0.53 | 0.951 |

**Table S7**: **Summary of GLMM results for predictors of interest in Experiment 3.** The reference condition is control and the reference image carcass. Significant effects are highlighted in bold. Significance levels: ***$p$ < 0.001, **$p$ < 0.01, *$p$ < 0.05.

| Model | Predictor | Estimate ($\beta$) | SE | $z$ | $p$-value |
|---|---|---|---|---|---|
| [M3.3] Latency | (Intercept) | 8.251 | 0.064 | 129.24 | $< 2e - 16 * **$ |
| | Condition:Test | 0.014 | 0.022 | 0.66 | 0.506 |
| | Image:Snake | 0.003 | 0.022 | 0.12 | 0.903 |
| | Exposure | -0.009 | 0.011 | -0.75 | 0.452 |
| | Test*Snake | -0.025 | 0.030 | -0.81 | 0.417 |
| | Test*Exposure | 0.014 | 0.016 | 0.91 | 0.362 |
| | Snake*Exposure | 0.005 | 0.016 | 0.33 | 0.744 |
| | **Test*Snake*Exposure** | **-0.057** | **0.022** | **-2.59** | **< 0.01**** |

**Table S8**: **Estimated marginal means and pairwise comparisons of latency across image types (carcass vs. snake) and conditions (test vs. control) in Experiment 3.** Codes: C = Carcass; Ctrl = Control; S = Snake; T = Test. Comparisons of interest were selected and adjusted using the Bonferroni method. Significant effects are highlighted in bold. Significance levels: ***$p$ < 0.001, **$p$ < 0.01, *$p$ < 0.05.

| Exposure | Condition | EMM (95%) | Contrast | $\beta$ | SE | $z$ | $p$-value |
|---|---|---|---|---|---|---|---|
| Min | C*Ctrl | 8.26 [8.13, 8.39] | - | - | - | - | - |
| | C*T | 8.26 [8.13, 8.39] | C*Ctrl - S*Ctrl | 0.005 | 0.033 | 0.16 | 1.000 |
| | S*Ctrl | 8.26 [8.13, 8.39] | C*T - S*T | -0.059 | 0.033 | -1.81 | 0.279 |
| | S*T | 8.32 [8.19, 8.45] | C*Ctrl - C*T | 0.008 | 0.033 | 0.24 | 1.000 |
| | | | S*Ctrl - S*T | -0.057 | 0.032 | -1.76 | 0.316 |
| Mid | C*Ctrl | 8.25 [8.13, 8.38] | - | - | - | - | - |
| | C*T | 8.27 [8.14, 8.39] | C*Ctrl - S*Ctrl | -0.003 | 0.022 | -0.12 | 1.000 |
| | S*Ctrl | 8.25 [8.13, 8.38] | C*T - S*T | 0.022 | 0.022 | 0.99 | 1.000 |
| | S*T | 8.24 [8.12, 8.37] | C*Ctrl - C*T | -0.014 | 0.022 | -0.67 | 1.000 |
| | | | S*Ctrl - S*T | 0.010 | 0.022 | 0.46 | 1.000 |
| High | C*Ctrl | 8.24 [8.11, 8.37] | - | - | - | - | - |
| | C*T | 8.27 [8.15, 8.40] | C*Ctrl - S*Ctrl | -0.011 | 0.033 | -0.32 | 1.000 |
| | S*Ctrl | 8.25 [8.12, 8.38] | **C*T - S*T** | **0.102** | **0.032** | **3.17** | **0.006**** |
| | S*T | 8.17 [8.04, 8.30] | C*Ctrl - C*T | -0.037 | 0.032 | -1.14 | 1.000 |
| | | | S*Ctrl - S*T | 0.076 | 0.033 | 2.28 | 0.090 |

**Table S9**: **Summary of GLMM results for predictors of interest in Experiment 4.** The reference condition is carcass. Significant effects are highlighted in bold. Significance levels: ***$p$ < 0.001, **$p$ < 0.01, *$p$ < 0.05.

| Model | Predictor | Estimate ($\beta$) | SE | $z$ | $p$-value |
|---|---|---|---|---|---|
| [M4.1] First fixation location | (Intercept) | 1.411 | 0.244 | 5.78 | 7.6e-9*** |
| | Snake | 0.001 | 0.237 | 0.01 | 0.995 |
| | **Tortoise** | **-0.808** | **0.299** | **-2.70** | **0.007**** |
| | **Pigeon** | **-1.387** | **0.298** | **-4.66** | **< 0.001***** |
| | **Deer** | **-0.854** | **0.330** | **-2.59** | **< 0.01**** |
| [M4.2] Proportion of fixations | (Intercept) | 0.522 | 0.106 | 4.94 | 7.9e-7*** |
| | Snake | 0.032 | 0.061 | 0.526 | 0.599 |
| | **Tortoise** | **-0.193** | **0.096** | **-2.01** | **0.044*** |
| | **Pigeon** | **-0.453** | **0.090** | **-5.05** | **< 0.001***** |
| | **Deer** | **-0.658** | **0.095** | **-6.90** | **< 0.001***** |
| [M4.3] Proportion of time looking | (Intercept) | 0.723 | 0.131 | 5.54 | 3.1e-8*** |
| | **Snake** | **0.157** | **0.074** | **2.12** | **0.034*** |
| | **Tortoise** | **-0.247** | **0.115** | **-2.15** | **0.032*** |
| | **Pigeon** | **-0.565** | **0.108** | **-5.25** | **< 0.001***** |
| | **Deer** | **-0.787** | **0.115** | **-6.83** | **< 0.001***** |

**Table S10**: **Estimated marginal means and pairwise comparisons of visual attention across image conditions in Experiment 4.** Visual attention is represented by the first fixation location, proportion of fixations, and proportion of time spent looking. Image codes: C = Carcass; D = Deer; P = Pigeon; S = Snake; T = Tortoise. Pairwise comparisons are Tukey-adjusted. Significance levels: ***$p$ < 0.001, **$p$ < 0.01, *$p$ < 0.05.

| Model | Cond | EMM (95% CI) | Contrast | $\beta$ | SE | $z$ | $p$-value |
|---|---|---|---|---|---|---|---|
| [M4.1] First fixation | C | 1.41 [0.93, 1.89] | - | - | - | - | - |
| | D | 0.56 [-0.10, 1.21] | C - D | 0.854 | 0.330 | 2.59 | 0.097 |
| | P | 0.02 [-0.57, 0.62] | **C - P** | **1.387** | **0.298** | **4.66** | **< 0.001*****|
| | S | 1.41 [0.92, 1.90] | C - S | -0.001 | 0.237 | -0.01 | 1.000 |
| | T | 0.60 [0.00, 1.20] | C - T | 0.808 | 0.299 | 2.70 | 0.069 |
| | | | S - D | 0.856 | 0.333 | 2.57 | 0.101 |
| | | | **S - P** | **1.388** | **0.303** | **4.58** | **< 0.001*****|
| | | | S - T | 0.809 | 0.305 | 2.66 | 0.079 |
| | | | D - P | 0.533 | 0.363 | 1.47 | 1.000 |
| | | | D - T | -0.046 | 0.366 | -0.13 | 1.000 |
| | | | P - T | -0.579 | 0.336 | -1.72 | 0.853 |
| [M4.2] Prop fixations | C | 0.52 [0.31, 0.73] | - | - | - | - | - |
| | D | -0.14 [-0.39, 0.12] | **C - D** | **0.658** | **0.095** | **6.90** | **< 0.001*****|
| | P | 0.07 [-0.18, 0.31] | **C - P** | **0.453** | **0.090** | **5.05** | **< 0.001*****|
| | S | 0.55 [0.34, 0.76] | C - S | -0.032 | 0.061 | -0.53 | 1.000 |
| | T | 0.33 [0.08, 0.58] | C - T | 0.193 | 0.096 | 2.01 | 0.260 |
| | | | **S - D** | **0.690** | **0.096** | **7.18** | **< 0.001*****|
| | | | **S - P** | **0.485** | **0.091** | **5.31** | **< 0.001*****|
| | | | S - T | 0.225 | 0.097 | 2.31 | 0.142 |
| | | | D - P | -0.205 | 0.103 | -1.99 | 0.272 |
| | | | **D - T** | **-0.466** | **0.109** | **-4.28** | **< 0.001*****|
| | | | P - T | -0.261 | 0.105 | -2.49 | 0.092 |
| [M4.3] Prop time | C | 0.72 [0.47, 0.98] | - | - | - | - | - |
| | D | -0.06 [-0.37, 0.25] | **C - D** | **0.787** | **0.115** | **6.83** | **< 0.001*****|
| | P | 0.16 [-0.14, 0.46] | **C - P** | **0.565** | **0.108** | **5.25** | **< 0.001*****|
| | S | 0.88 [0.62, 1.14] | C - S | -0.157 | 0.074 | -2.12 | 0.210 |
| | T | 0.48 [0.17, 0.79] | C - T | 0.247 | 0.115 | 2.15 | 0.319 |
| | | | **S - D** | **0.944** | **0.117** | **8.08** | **< 0.001*****|
| | | | **S - P** | **0.722** | **0.110** | **6.55** | **< 0.001*****|
| | | | **S - T** | **0.404** | **0.118** | **3.44** | **0.005****|
| | | | D - P | -0.222 | 0.125 | -1.78 | 0.386 |
| | | | **D - T** | **-0.539** | **0.132** | **-4.10** | **< 0.001*****|
| | | | P - T | -0.318 | 0.126 | -2.53 | 0.085 |

**Movie S1. Chimpanzee Ai performing the number ordering task under the cadaverine condition (Session 1, Experiment 2).** Low concentrations of cadaverine were released beneath the screen while chimpanzees completed the task.

**Movie S2. Chimpanzee Cleo performing the number ordering task under the snake condition (Session 2, Experiment 3).** Snake images were displayed full-screen every five trials over 50 trials. Chimpanzees performed faster in this condition compared to the carcass condition.

**Movie S3. Chimpanzee Cleo performing the number ordering task under the carcass condition (Session 1, Experiment 3).** Images of heterospecific animal carcasses were displayed full-screen every five trials over 50 trials. Chimpanzees performed slower in this condition compared to the snake condition.

**Movie S4. Chimpanzee Pendesa performing the number ordering task under the disease-associated invertebrate condition (Session 2, Experiment 1).** Images of invertebrates were displayed full-screen every five trials, following nine initial warm-up trials. Pendesa's number set was reduced from 9 to 6 to accommodate her motivation and capacities. Here, she turns away from the screen when presented with picture 3.

**Movie S5. Chimpanzee Cleo performing the eye-tracking task under the tortoise condition (Experiment 4).** The session began with calibration using small moving icons paired with sounds, followed by a corner calibration. Each trial then consisted of a 5-second image presentation, after which corner calibration was repeated to decentralize gaze. This sequence was repeated until 10 images had been presented. Gaze plots illustrate fixations, with circle size reflecting fixation duration, and saccades (lines between circles).
